# NE52-QQ57 attenuates post-ischemia protein phosphorylation in the mouse brain

**DOI:** 10.64898/2026.10.09.754558

**Authors:** Shambhu Joshi, Youfang Zhou, Xiang-ming Zha

**Author notes:** Address correspondence to: Xiangming Zha, PhD Department of Neurosurgery Tulane University School of Medicine 1324 Tulane Ave New Orleans, LA 70124.

## Abstract

**Introduction:** Cerebral ischemia induces a reduction in brain pH. GPR4, a proton-sensing G protein-coupled receptor is highly expressed in vascular endothelial cells. Pharmacological inhibition of GPR4 with NE52-QQ57 (NEQQ) reduces ischemia-induced blood-brain barrier (BBB) disruption and brain injury. To facilitate targeting GPR4 in vivo, it is important to better define the effect of NEQQ on post-ischemia signaling and the pharmacokinetics of NEQQ *in vivo*.

**Methods:** We performed biochemical analysis following transient middle cerebral artery occlusion (tMCAO). We used Liquid Chromatography-Mass spectrometry (LC-MS/MS) to quantify the concentration of NEQQ in serum, brain, and peripheral tissues.

**Results:** NEQQ administration decreased the phosphorylation of PKA substrates, PKC substrates, ERK1/2, and CREB. LC-MS/MS analysis detected NEQQ in both ipsilateral and contralateral hemisphere of the brain at 5 h and 24 h after reperfusion, demonstrating that NEQQ crosses the BBB. In peripheral tissues, NEQQ exhibited highest level in kidney and lowest in the heart. In sham controls, average serum concentrations were ∼4 and 0.9 µg/g at 5 and 24 h, respectively. Following tMCAO, ipsilateral brain exhibited higher levels as compared to the contralateral side at 5 h while the difference diminished at 24 h.

**Conclusions:** NEQQ attenuated the ischemia-induced activation of multiple signaling pathways. NEQQ exhibited broad distribution in peripheral organs, and its brain level was sufficient for GPR4 inhibition at 24 h after injection. These findings support the use of NEQQ as a pharmacological inhibitor of GPR4-mediated signaling following ischemic stroke.

## Introduction

Stroke affects millions of people worldwide, has high mortality, and leads to long-term disability (1). Most strokes are ischemic and occur when an artery supplying the brain becomes suddenly blocked, restricting the blood flow to brain tissue (2). Ischemia leads to the acidification of brain parenchyma (3, 4), with pH lowering to the range of 6.5–6.0 (5–7). G protein-coupled receptor 4 (GPR4) is a proton-sensing G protein-coupled receptor that is highly expressed in vascular endothelial cells (8). An *in vitro* study has further demonstrated that extracellular acidosis induces GPR4 activation and resulted into significant increase in the paracellular gap formation and permeability in human umbilical vein endothelial cells (8). Our recent data showed that GPR4 in the brain contributes to ischemia-induced blood-brain barrier (BBB) dysfunction.

NEQQ is a pyrazolopyrimidine derivative and a highly selective antagonist of GPR4 (9, 10). Pharmacological inhibition of GPR4 with NEQQ was reported to mitigate blood vessel permeability, tissue edema, and leukocyte infiltration in the mouse model of acute hindlimb ischemia-reperfusion (8). In preclinical models, NEQQ has also demonstrated anti-inflammatory property including reduction in arthritic inflammation, hyperalgesia, and colitis (9, 11). We have data to show that GPR4 inhibition with NEQQ protected against acidosis-induced brain microvascular endothelial cell (BMEC) junctional dysfunction and reduced ischemia-induced BBB leakage. These findings support a promising therapeutic potential of NEQQ. However, the effects of NEQQ on ischemia-induced GPR4 signaling in vivo and its pharmacokinetics following systemic administration has not been well studied.

In this study, we started by asking whether NEQQ alters post-ischemia signaling in vivo. Next, we characterized the biodistribution of NEQQ in mice. To define time-dependent changes in brain: serum ratio, we determined NEQQ levels in the brain and serum at 5 and 24 h following ischemia.

## Materials and Methods

### Mice

To analyze NEQQ level and signaling in vivo, we used GPR4 +/+ mice on both C57BL6/N and C57BL6/J background. Mice were generated through in-house breeding. Male mice aged 12-37 weeks and female mice aged 16-28 weeks were used in this study. All mice were housed under standard laboratory conditions with free access to food and water. All experimental procedures were conducted in accordance with the National Institutes of Health Guide for the Care and Use of Laboratory Animals and were approved by the Tulane University Institutional Animal Care and Use Committee.

### Reagents and antibodies

NEQQ was purchased from MedChemExpress (Catalog # HY-101784) and Cayman Chemical (Catalog # 40647). Primary antibodies used for Western blotting (WB) includes: Rb Phospho-PKA Substrate (RRXS*/T*) (100G7E, Cell Signaling Technology, Catalog # 9624, 1:2000); Rb Phospho-(Ser) PKC Substrate (Cell Signaling Technology, Catalog # 2261, 1:2000), Ms beta tubulin, E7 (Developmental Studies Hybridoma Bank, 1:10,000, RRID: AB_528499), Rb Phospho-ERK1/2 (Upstate, Catalog # 07-467, 1:1000), Ms ERK1/2 (Invitrogen, Catalog # MA5-15227, 1:1000, RRID AB_10980663), Rb Phospho-CREB (Ser 133) (87G3, Cell Signaling Technology, Catalog # 9198, 1:1000), Ms CREB (86B10) (Cell Signaling Technology, Catalog # 9104, 1:1000). Secondary antibodies used for WB include donkey anti rabbit IRdye 800 (1:10 K, (Li-Cor Biosciences, Catalog # 926– 32,213) and donkey anti-mouse IRdye 680 (1:10 K, Li-Cor Biosciences, Catalog # 926–68,022).

### Transient middle cerebral artery occlusion (tMCAO)

Transient focal cerebral ischemia was induced by suture occlusion of the middle cerebral artery (MCAO) using the Longa model as described previously (12–14). Briefly, male (28 ± 4 g) and female (23 ± 3 g) mice were randomly allocated to experimental groups. Mice were anesthetized with 1.5–2% isoflurane in a 70% N₂O/30% O₂ mixture, and body temperature was maintained at 37 ± 0.5 °C using a feedback-controlled heating pad throughout the procedure. Cerebral blood flow (CBF) was continuously monitored using laser Doppler flowmetry (MoorVMS-LDF2), with the probe fixed to the skull using cyanoacrylate adhesive. A silicone-coated monofilament (Doccol Corp.) was inserted into the external carotid artery (ECA) and advanced into the internal carotid artery (ICA) to occlude the middle cerebral artery (MCA). Filament size was selected based on body weight. We used suture #702023 for mice weighing < 28 g) and suture #702223 for mice weighing 28–34 g. After 60 min of MCA occlusion, the filament was withdrawn to allow reperfusion. For CBF analysis, mean flow values were recorded at baseline, during MCA occlusion, and after reperfusion (average of a 1–2 min peak value). Mice were included for analysis only when CBF was reduced to 5–20% of baseline during occlusion and recovered to ≥55% following reperfusion.

### Western blotting

The ipsilateral and contralateral hemispheres of the mouse brain tissues were collected separately and homogenized in lysis buffer consisting of 1% Triton, 0.4% SDS, protease inhibitors (Roche) and phosphatase inhibitors (Roche) diluted with 1X PBS. The tissue lysates were then incubated for 10 min on ice, and subsequently sonicated. The lysates were centrifuged at 15,000 x g for 10 min at 4°C and the resulting supernatants were collected as a total protein lysate. Protein concentration was determined using Bio-Rad Protein Assay kit and equal amounts of total protein (5 to 20 μg) were separated by SDS-PAGE on 8-12% polyacrylamide gels at 110V for 90 minutes. The gels were subsequently transferred onto nitrocellulose membrane (Bio-Rad) at 100 mA for 16 h. Membranes were blocked for 1 h with the blocking buffer containing 0.1% casein in 0.2 x PBS pH 7.2-7.6 for 1 h. The membranes were then incubated overnight at 4°C with following primary antibodies: Rb Phospho-PKA Substrate (RRXS*/T*) (100G7E, Cell Signaling Technology, Catalog # 9624, 1:2000); Rb Phospho-(Ser) PKC Substrate (Cell Signaling Technology, Catalog # 2261, 1:2000), Ms beta tubulin, E7 (Developmental Studies Hybridoma Bank, 1:10,000, RRID: AB_528499), Rb Phospho-ERK1/2 (Upstate, Catalog # 07-467, 1:1000), Ms ERK1/2 (Invitrogen, Catalog # MA5-15227, 1:1000, RRID AB_10980663), Rb Phospho-CREB (Ser 133) (87G3, Cell Signaling Technology, Catalog # 9198, 1:1000), Ms CREB (86B10) (Cell Signaling Technology, Catalog # 9104, 1:1000). After washing with PBST (PBS containing 0.05% Tween-20), the membranes were incubated at room temperature for 1 hour with secondary antibodies. Following additional washing the membranes in PBST, the blots were imaged using ChemiDoc MP Imaging System (Bio-Rad) according to manufacturer’s instructions. Image J was used for densitometric analysis of imaged bands.

### Drug delivery, tissue collection and LC-MS/MS analysis

Mice underwent 60 min transient middle cerebral artery occlusion (tMCAO), followed by reperfusion. After 2 h of reperfusion, NEQQ was administered intraperitoneally (i.p.) at 30 mg/kg body weight. Samples were collected at 5 and 24 h after reperfusion. Blood was collected from the heart immediately before transcardial perfusion with ice-cold saline or without perfusion. Serum was isolated and stored at -80°C until analysis. In perfused mice, the ipsilateral and contralateral brain hemispheres were collected separately following saline perfusion. In non-perfused mice, the ipsilateral and contralateral hemispheres were collected separately without prior perfusion. Peripheral organs, including the heart, lungs, liver, spleen and kidneys were also collected from both perfused and non-perfused mice. All tissue samples were immediately frozen and stored at -80°C until LC-MS/MS analysis for NEQQ quantification.

For LC-MS/MS analysis of NEQQ, the service was provided by WuXi AppTec. Briefly, concentration of NEQQ in mouse serum and tissue samples including brain, the heart, lungs, liver, spleen and kidneys were quantified using a Triple Quad 6500+ mass spectrometer equipped with an electrospray ionization (ESI) source operated in positive-ion mode. The chromatographic separation was performed on a Waters ACQUITY UPLC BEH C18 column (2.1 × 50 mm, 1.7 µm) using 0.1% formic acid in water (Mobile phase A) and 0.1% formic acid in acetonitrile (Mobile phase B) at a flow rate of 0.6 mL/min. Quantification was done using multiple reaction monitoring (MRM). NEQQ was detected as the protonated molecular ion [M+H]+ at m/z 417.2>292.2. Labetalol and tolbutamide were used as internal standards with MRM transitions of [M+H]+ m/z at 329.2>162.1 and [M+H]+ at m/z 271.1>155.3 respectively each prepared at 100 ng/mL in acetonitrile. Calibration curves for the NEQQ were generated over the range of 1-3000 ng/mL for mouse brain and 5-in-1 tissue matrix samples and over the range of 1-3000 ng/mL in serum samples. Data was acquired and analyzed using the instrument associated mass spectrometry software.

For tissue samples, tissues were homogenized in a ratio of 1:5 (w/v) using homogenate buffer (1x PBS: MeOH, v/v = 2:1). Serum or tissue homogenate (20 μL) was protein-precipitated with 200 µL internal standard (IS) solution (100 ng/mL Labetalol & 100 ng/mL Tolbutamide in acetonitrile), the mixture was vortex-mixed well and centrifuged at 4000 rpm (tissue) or 3900 rpm (serum) for 10 min at 4 ℃. Subsequently, an aliquot of 100 μL supernatant was transferred to a sample plate, mixed with 100 μL water, and shaken at 800 rpm for 5 min. An aliquot of 0.5 µL supernatant was then injected for LC-MS/MS analysis.

### Statistical Analysis

Statistics were performed in GraphPad Prism and Microsoft Excel. Data normality was assessed to determine the downstream tests to use. For comparing 2 groups, depending on the normality test result, the 2-tailed *t* test, the Mann-Whitney *U* test was used. Data are presented as mean ± SEM. Differences were considered significant if *p* < 0.05.

## RESULTS

### NEQQ attenuates the post-ischemia phosphorylation of multiple pathways

In our recent study, NEQQ delivered at reperfusion reduced tMCAO-induced brain infarct size at 24 hours. This raised the question of whether NEQQ IP at reperfusion is sufficient to attenuate ischemia-induced signaling at the 24-hour time point. To answer this question, we performed 60 min tMCAO in GPR4+/+ mice, then administered vehicle or NEQQ (30 mg/kg) via IP at 2 h of reperfusion, isolated brain tissue at ∼24 h, and analyzed several ischemia-induced signaling pathways described by previous literature (15–23). As expected, ischemia increased the phosphorylation of PKA substrates, PKC substrates, Erk1/2, and CREB (Figure 1). NEQQ treatment attenuated the phosphorylation of these targets in both the ipsilateral and contralateral brain regions compared with vehicle-treated mice (Fig 1).

**Figure 1:**
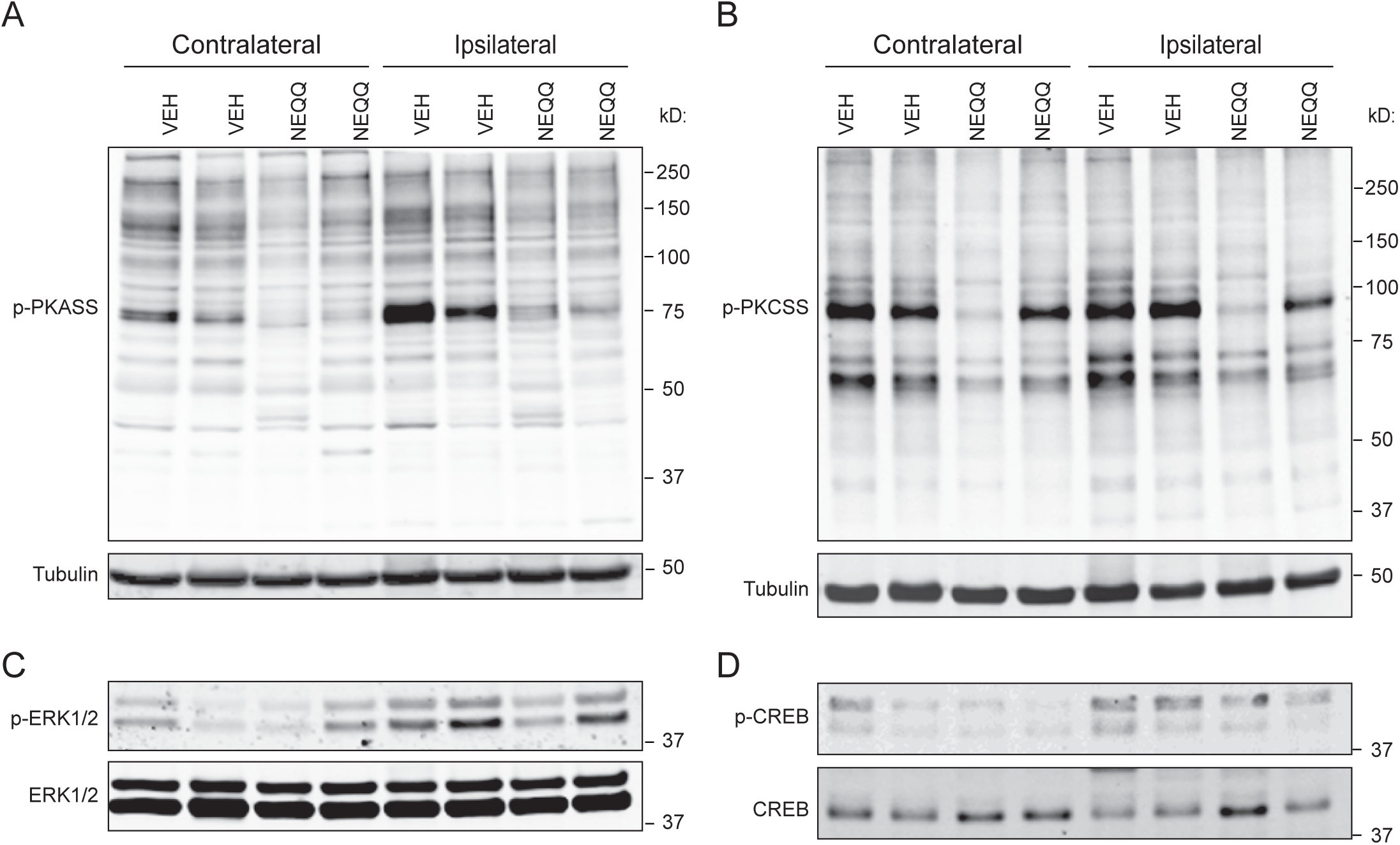
NEQQ attenuates the ischemia-induced signaling. Representative Western blot images showing the effects of NEQQ treatment on ischemia-induced signaling in the brain. At 2-hour following the 60 min tMCAO, we administered vehicle or NEQQ at 30 mg/kg, i.p., collected tissue at 24 hour and performed Western blotting analysis. (A) NEQQ reduced phosphorylation of PKA substrates in both the ipsilateral and contralateral brain regions. (B) NEQQ reduced phosphorylation of PKC substrates in both the ipsilateral and contralateral brain regions. (C) NEQQ attenuated ischemia-induced ERK1/2 phosphorylation in the ipsilateral brain region. (D) NEQQ attenuated ischemia-induced CREB phosphorylation in the ipsilateral brain region.

### Detection of NEQQ in mouse brain tissue using LC-MS/MS

The above analysis showed that NEQQ is sufficient to impact post-ischemia signaling at up to 24 h after reperfusion. To better define the in vivo kinetics of NEQQ, we performed LC-MS/MS analysis (LC-MS/MS service provided by Wuxi AppTec). First, we validated the extraction and detection of NEQQ in brain and serum samples. To mouse brain tissue lysate or serum, we added different concentrations of NEQQ. Following extraction, we employed LC-MS/MS to detect and quantify NEQQ in tissue samples. Figure 2 shows representative multiple-reaction monitoring (MRM) chromatograms for detection of NEQQ and internal standard (IS) in mouse brain samples. No distinctive NEQQ signal was observed in the blank brain sample, whereas NEQQ were detected in standards, ranging from 1 ng/mL to 3000 ng/mL (Fig 2A). From these, we generated a standard curve for calibrating the level of NEQQ in experimental samples (Fig 2B).

**Figure 2:**
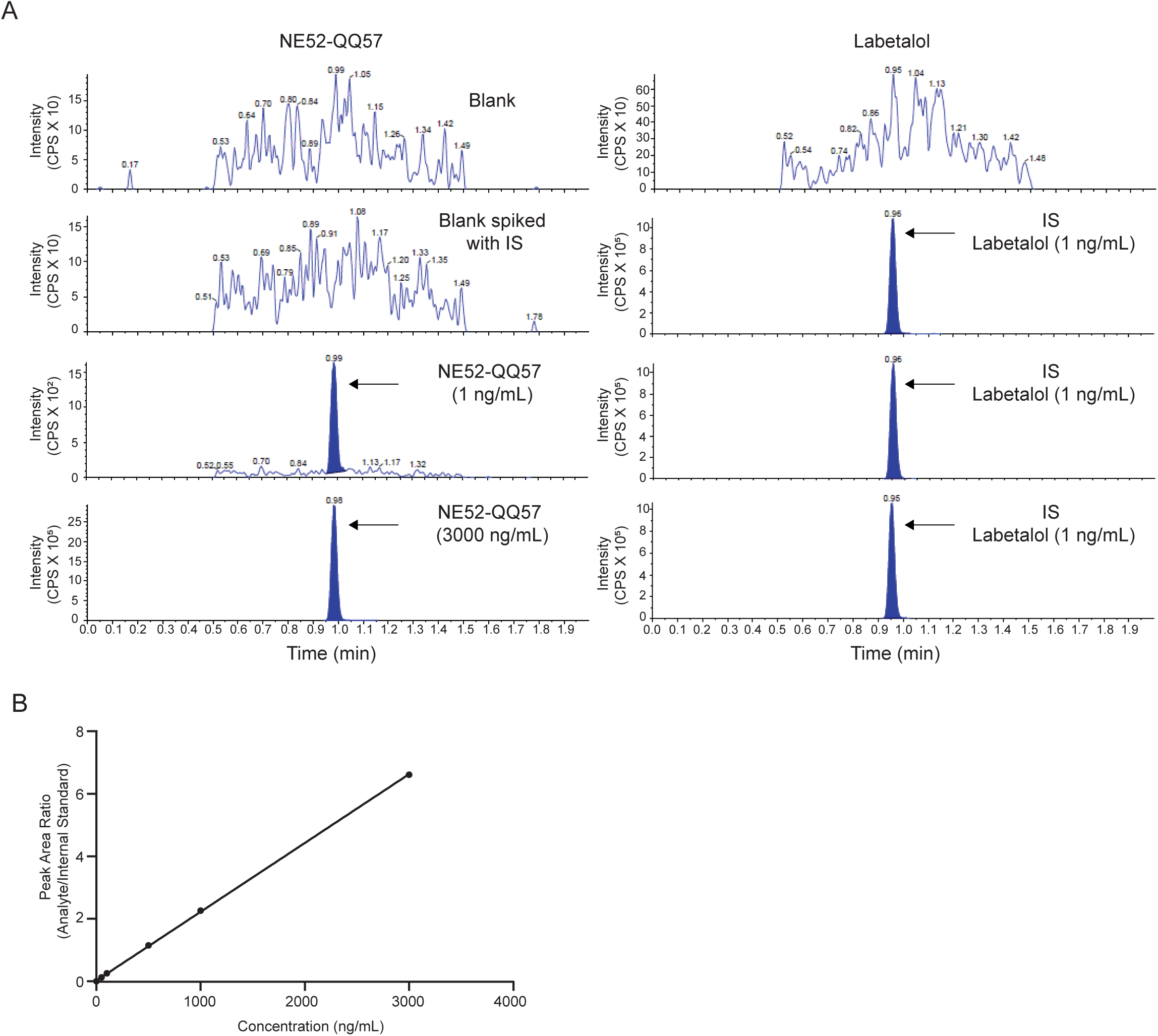
NEQQ concentration was quantified by LC-MS method. (A) Representative multiple reaction monitoring (MRM) chromatograms of NEQQ and the internal standard (IS) obtained from a blank mouse brain without analyte or IS, a blank mouse brain containing the IS (zero sample), a blank mouse brain spiked with NEQQ at the lower limit of quantification (LLOQ;1 ng/mL or 6 ng/g) along with the IS and a blank mouse brain spiked with NEQQ at upper limit of quantification (ULOQ; 3000 ng/mL or 18000 ng/g) along with IS (B) Standard curve used to quantify NEQQ concentrations in experimental samples.

To determine NEQQ level following delivery in vivo, we administered NEQQ (30 mg/kg, i.p.) in sham operated or tMCAO mice, collected tissue at 5 and 24 h after reperfusion following tMCAO (Fig 3A). NEQQ was readily detected in serum and brain tissues from both sham-operated and tMCAO mice. In sham operated mice, serum NEQQ concentrations were 3877.5 and 909 ng/mL at 5 and 24 h, respectively. In tMCAO mice, serum NEQQ concentrations were 8570 and 1173.75 ng/mL at 5 and 24 h, respectively (Fig 3B, C).

**Figure 3:**
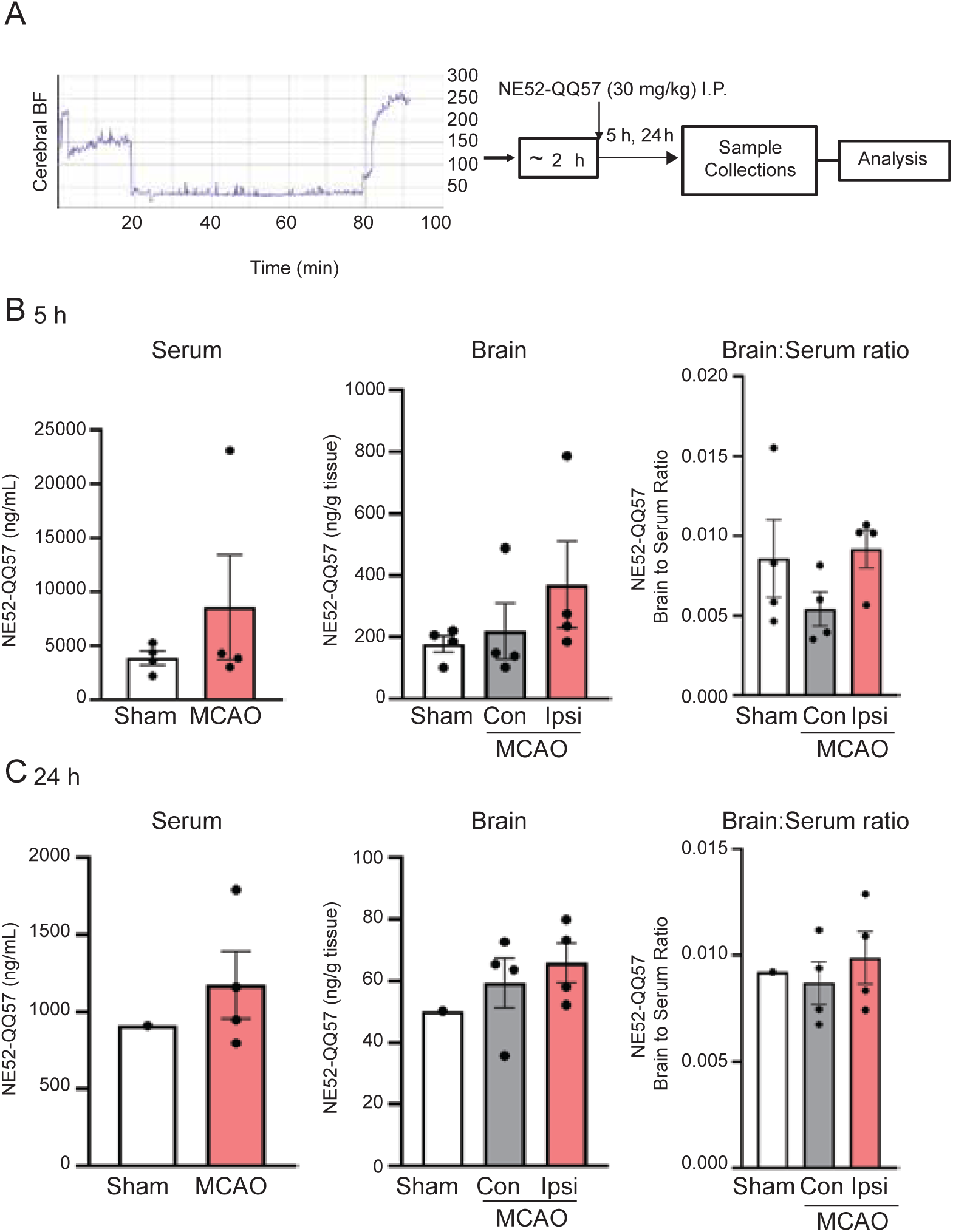
Biodistribution of NEQQ in serum and brain tissue following tMCAO. (A) Representative cerebral flow (CBF) during tMCAO and experimental timeline. (B) Serum NEQQ concentration, brain tissue NEQQ concentration and brain to serum NEQQ concentration ratio after 5 h of reperfusion. (B) Serum NEQQ concentration, brain tissue NEQQ concentration and brain to serum NEQQ concentration ratio after 24 h of reperfusion in sham-operated and tMCAO mice. NEQQ concentrations in brain tissue were measured separately for the ipsilateral and contralateral hemispheres. NEQQ was administered at 30 mg/kg, i.p., 2 h after initiation of reperfusion. NEQQ concentrations in serum and brain tissue were quantified by LC-MS/MS.

We next analyzed the level of NEQQ in the brain. In sham-operated mice, NEQQ concentrations in the brain were 178.05 ng/g at 5 h and 50.2 ng/g at 24 h. At 5h, NEQQ concentrations were higher in the ipsilateral hemisphere (370.5 ng/g) than in the contralateral hemisphere (219.45 ng/g). At 24 h, NEQQ concentration in the brain was 65.81 and 59.33 ng/g, respectively in the ipsilateral and contralateral hemisphere (Fig 3B, C). These findings demonstrate that NEQQ reaches the brain following systemic administration, with greater concentration in the ipsilateral hemisphere after tMCAO.

To further characterize brain exposure to NEQQ following systemic administration, we calculated the brain to serum concentration ratio. The brain to serum concentration ratio was less ∼ 1% at both 5 and 24 h after reperfusion, indicating that NEQQ concentrations in the brain were substantially lower than those in serum (Fig 3B, C). Despite the relatively low brain to serum ratio, NEQQ was readily detected in both ipsilateral and contralateral regions following systemic administration, demonstrating measurable brain exposure to the NEQQ.

To determine tissue distribution, we collected tissue from major organs and performed similar analysis. At 5 h after reperfusion, NEQQ exhibited the highest concentration in the kidney (13260 ng/g). This is followed by spleen (9550 ng/g), lungs (7012 ng/g), liver (5888 ng/g) and heart (2260 ng/g) (Fig 4).

**Figure 4:**
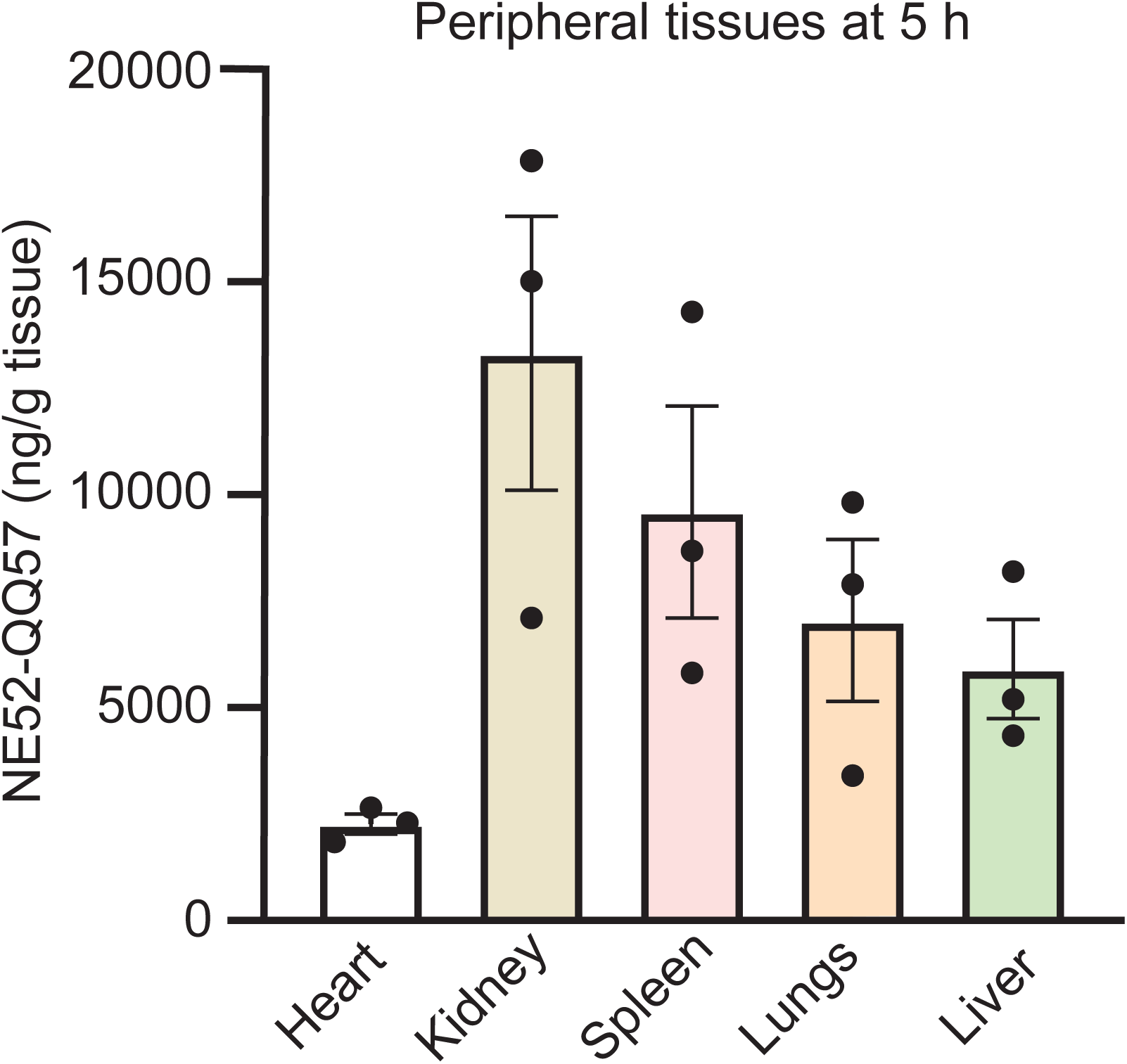
Biodistribution of NEQQ in peripheral tissues at 5. **h.** NEQQ concentrations in peripheral tissues 5 h after reperfusion, with the highest and lowest concentrations observed in the kidney and heart, respectively. NEQQ was administered at 30 mg/kg, i.p., 2 h after initiation of reperfusion. NEQQ concentrations were quantified by LC-MS/MS.

## Discussions

In the present study, we characterized the *in vivo* tissue distribution and pharmacodynamic analysis of NEQQ following systemic administration in the post stroke condition. The LC-MS/MS result showed that 0.026 µM of NEQQ in the brain remained at 24 h after a single injection of NEQQ at 2 h of reperfusion. This result is consistent with the persistent effect of NEQQ on post-ischemia signaling at this time point. These findings provide new insights into pharmacokinetic properties of NEQQ and how it regulates post-ischemia signaling.

NEQQ is orally active (9, 24). Previous studies have evaluated NEQQ as a GPR4 antagonist using both oral and intraperitoneal administration (9–11, 24). Following stroke surgery, oral gavage is challenging for mice in the initial days. For this reason, we delivered NEQQ via IP, and detected NEQQ in serum, brain, and all peripheral tissues. One technical note is that IP administration bypasses gastrointestinal metabolism and increases bioavailability as compared with oral gavage (25). In both sham and stroke mice, we detected NEQQ in brain tissue. This is consistent with previous report which shows that NEQQ passes BBB (10, 26). Moreover, our data showed that NEQQ concentration was higher in the ipsilateral hemisphere (370.50 ng/g) than in the contralateral hemisphere (219.45 ng/g) at 5 h after reperfusion. This regional difference is consistent with the well-established increase in cerebrovascular permeability that initiates after cerebral ischemia with the initial phase of BBB leakage occurs at around 3 to 6 hours following ischemia-reperfusion (27–29).

Increased vascular permeability may therefore contribute to enhanced accumulation of NEQQ in the ipsilateral hemisphere. Nevertheless, the detection of NEQQ in the contralateral and sham-operated brains indicates that its CNS distribution does not depend solely on ischemia-induced BBB disruption. Together, these findings support CNS exposure to NEQQ under both physiological and post-stroke conditions with potentially enhanced exposure in the ischemic hemisphere during the early phase of reperfusion.

NEQQ exposure was consistently higher at 5 h compared to 24 h in serum and brain tissue, indicating a decline in systemic and tissue exposure over time. This temporal pattern is compatible with drug elimination and/or redistribution between tissues; however, our experimental design and sampling does not allow us to specifically determine the specific elimination pathway or pharmacokinetic parameters such as clearance and half-life. NEQQ was also detected in all major organs examined, including the liver, kidney, spleen, lungs and heart. At 5 h, the kidney showed the highest concentration (13260 ng/g) and this is followed by spleen, lungs, liver and heart (2260 ng/g) indicating substantial peripheral distribution. The pronounced renal accumulation is consistent with a potential role of the kidney in drug disposition, although direct evidence of renal elimination would require measurement of urinary excretion and/or additional pharmacokinetic analyses.

Cerebral ischemia is accompanied by tissue acidosis, creating a pathophysiological environment in which proton-sensing receptors such as GPR4 become activated (3, 4) and subsequently activates downstream signaling through Gs/cAMP, Gq/phospholipase C (PLC), and G13/Rho pathways (30–33). We therefore examined selected downstream signaling readouts to determine whether systemic NEQQ administration was associated with modulation of GPR4-associated signaling in the ischemic brain. Our data on signaling pathway demonstrates that NEQQ when administered at 30 mg/kg, i.p. decreases the phosphorylation of PKA and PKC substrates in both ipsilateral and contralateral brain hemispheres suggesting that these brain hemispheres are exposed to the drug at pharmacologically active level. However, these signaling changes should not be interpreted as direct evidence of exclusive GPR4 inhibition, because the measured pathways can also be regulated by other signaling mechanisms during ischemic injury. For example, the expression of GPR68, another protein sensing G protein coupled receptor expression at mRNA level is increased following ischemia (14) which could also contribute to the observed effects on the signaling. Inhibition of GPR4 mediated signaling pathway may represent a therapeutically relevant mechanism for mitigating the signaling associated with ischemic vascular dysfunction and a better understanding of acid signaling will have important clinical implications for designing better treatment for stroke. The observed attenuation of phosphorylated ERK1/2 and CREB signaling is of particular interest because these pathways have been implicated in the process relevant to ischemic injury, including inflammation, cell survival and death, and BBB dysfunction (16–19). Although the present study does not establish that each of these signaling changes is mediated exclusively through GPR4, the combined pharmacokinetic and pharmacodynamic findings provide functional support for CNS exposure and modulation of GPR4-associated signaling by NEQQ following systemic administration. Thus, the present findings provide a pharmacological framework for further investigating the contribution of GPR4-dependent acid signaling to ischemic vascular dysfunction and stroke pathology.

The detection of NEQQ at considerable amount in the peripheral tissues also raises the possibility that its pharmacological effects extend beyond the brain. This consideration is particularly relevant to organs such as kidneys, where relatively high exposure was observed and where GPR4 plays an important role to maintain the acid-base balance (34). Further studies directly examining GPR4-associated signaling in peripheral tissues will be necessary to determine whether systemic NEQQ exposure exerts pharmacodynamic effects beyond CNS.

Collectively, these findings establish a link between systemic administration, tissue exposure and pharmacodynamic activity of NEQQ in the post-stroke brain. The ability of NEQQ to reach the CNS and modulate signaling associated with GPR4 provides a pharmacological basis for further investigating GPR4 as a therapeutic target in ischemic stroke. Further studies incorporating comprehensive pharmacokinetic profiling, metabolite characterization and direct assessment of downstream targets will be important for defining the exposure-pharmacodynamic response relationship and determining how NEQQ distribution relates to its therapeutic effects after ischemic stroke.

## Author Contribution

SJ and YZ performed transient middle cerebral artery occlusion (tMCAO) surgery and collected samples for analysis. SJ performed Western blot analysis, analyzed LC-MS/MS result, and wrote the original draft. XMZ conceived the idea, supervised, and edited the manuscript. All authors reviewed the final manuscript.

## Acknowledgement

We thank Angela Barattini and Usman Ali for technical assistance, Wenyan Sun for advice on MCAO surgery, and Yingjun (John) Ma at WuXi AppTec for assistance on the LC-MS/MS service.

